# Innate and adaptive immunity to transfused allogenic red blood cells in mice requires MyD88

**DOI:** 10.1101/2021.08.24.457332

**Authors:** Arielle Soldatenko, Laura R. Hoyt, Lan Xu, Samuele Calabro, Steven M. Lewis, Antonia E. Gallman, Krystalyn E. Hudson, Sean R. Stowell, Chance J. Luckey, James C. Zimring, Dong Liu, Manjula Santhanakrishnan, Jeanne E. Hendrickson, Stephanie C. Eisenbarth

## Abstract

Red blood cell (RBC) transfusion therapy is essential for the survival of patients with hematological disorders such as sickle cell anemia. A potentially fatal complication of transfusion is development of non-ABO alloantibodies to polymorphic RBC antigens, yet mechanisms of alloantibody formation remain unclear. Human and mouse RBCs acquire a “storage lesion” prior to transfusion, which in mice contributes to immunogenicity. We previously reported that mouse splenic dendritic cells (DCs) are required for RBC alloimmunization and are activated by sterile and leukoreduced mouse RBCs after storage. Yet how syngeneic RBCs activate innate immune pathways to induce DC activation is unknown. We now show that DC activation to transfused RBCs occurs regardless of alloantigen presence, suggesting that RBC damage induced during storage triggers innate immune receptors. We discovered an unexpected dependence of RBC alloimmunization on the Toll-like receptor (TLR) signaling adaptor molecule MyD88. TLRs are a class of pattern recognition receptors (PRRs) that regulate DC activation and signal through two adaptor molecules, MyD88 and TRIF. We show that the inflammatory cytokine response, DC activation, and the subsequent alloantibody response to transfused syngeneic RBCs require MyD88 but not TRIF, suggesting a restricted set of PRRs are responsible for sensing RBCs and triggering alloimmunization.

## Introduction

Chronic red blood cell (RBC) transfusion therapy is a mainstay of treatment for patients with hematological disorders and bone marrow failure syndromes such as sickle cell anemia, thalassemia major, and myelodysplastic syndrome. However, a major complication of RBC transfusion is the development of non-ABO alloantibodies (1). From infancy, humans constitutively produce IgM antibodies against A and B carbohydrate blood group antigens they lack (2). Unlike these natural antibodies, non-ABO alloantibodies target polymorphic minor antigens on donor RBCs following exposure via transfusion or pregnancy. RBC alloimmunization affects 3-10% of transfused patients and up to 50% of chronically transfused groups (3–5). As more than 12 million units of RBCs are transfused in the U.S. every year, the number of patients with alloantibodies is significant. RBC alloimmunization can result in hemolytic transfusion reactions, which are potentially fatal and can also delay the time to transfusion while compatible blood is sought (1, 6–9). Other than attempting to avoid transfusion altogether, there are limited therapeutic options to prevent RBC alloimmunization. Although extended antigen matching has been helpful, it is not feasible for many blood group antigens and for particular patients. Despite its clinical significance, the mechanism of alloantibody formation in response to RBC transfusion remains unclear.

In the United States, a typical sterile red cell unit for human transfusion is collected in anticoagulant, processed to remove leukocytes, and stored in special solutions to preserve RBC function at 1-6°C for up to 42 days at a hematocrit of approximately 60% (10). The average storage age of blood used for transfusions in the United States is estimated to be 18 days (11). During storage, RBCs can accumulate various types of damage, collectively termed “storage lesion”, which in mice has been shown to contribute to immunogenicity through an unknown mechanism (12). Our previous work and others have shown that transfusion of RBCs stored in this manner for as little as 12 days induces an inflammatory response in recipient mice and promotes subsequent alloantibody formation (13–15). Units of stored RBCs contain many potentially inflammatory components including iron, free heme, extracellular ATP, and modified lipids (14–15). Human studies suggest that transfusion in the presence of innate immune activation (through recipient inflammation) enhances RBC alloimmunization, however the specific role of storage itself on this effect is controversial (16–19). Identifying an effect of storage on alloimmunization in humans is difficult given the large number of alloantigens without routine follow up testing for alloantibody induction, potential use of immunosuppression in RBC recipients, the lack of alloantigen typing on the majority of RBC units, practical issues around RBC storage duration in blood bank inventories, and the lack of knowledge regarding recipient HLA capable of presenting polymorphic epitopes. Therefore, we used mouse models developed to study the T cell and B cell response to syngeneic murine RBCs containing model alloantigens to test the hypothesis that damage to the RBC itself during storage is essential for triggering innate immune pathways that induce non-ABO alloantibodies.

In order to mount an adaptive immune response, specifically naïve T and B cell priming, first the innate immune system must be activated. This involves activation of antigen-presenting cells, such as dendritic cells (DCs), through innate immune receptors termed pattern recognition receptors (PRRs); this is a crucial checkpoint in regulating T cell-dependent B cell antibody responses (20). Indeed, our previous work demonstrated that although transfused RBCs are consumed by and activate numerous antigen presenting cells in the spleen, only type 2 conventional dendritic cells (cDC2s) were required for alloantibodies (21), likely due to a defect in T cell priming (22). How RBC transfusion activates these DCs, however, is not known (13). Prior work from others has similarly shown that the same bridging channel cDC2s are responsible for T cell-dependent B cell responses to sheep RBCs; for these xenogeneic RBCS, DC activation is induced, at least in part, though species differences in CD47 (23, 24). As CD47 is not significantly altered by storage nor induces DC activation during allogenic transfusion (25, 26), we searched for other innate immune pathways operational during allogenic RBC transfusion. We hypothesized that the damage accumulated by RBCs during storage might stimulate an initial innate immune response through the ligation of one or several of PRRs on cDC2s.

PRRs sense both foreign motifs expressed by pathogens termed pathogen associated molecular patterns (PAMPs) and self-molecules released by or expressed on damaged host cells termed damage associated molecular patterns (DAMPs) (27). Our previous work demonstrated that potentially relevant NOD-like receptor (NLR) PRRs were not required for alloantibody formation, nor was the presence of the IL-1 family cytokines (IL-1 and IL-18) (13). Murine studies demonstrate that recipient exposure to various PAMPs such as Poly(I:C), CpG and influenza virus infection prior to transfusion of RBCs enhances alloimmunization to multiple alloantigens (28–30). These PAMPs are ligands for another family of PRRs, the Toll-like receptors (TLRs). Ten TLRs have been identified in humans and 13 in mice, which can be activated by both PAMPs and DAMPs and signal through the adapter proteins TRIF and/or MyD88 (31).

We used transgenic mice with RBCs expressing a model alloantigen that enabled tracking of the recipient immune response in order to test the hypothesis that stored RBCs activate TLRs on DCs to induce adaptive immunity (14,32). Here we show that MyD88, but not TRIF, is required for activation of splenic DCs by transfusion of stored RBCs. These results suggest that one or more TLRs dictate whether transfused RBCs induce an inflammatory response capable of promoting RBC alloimmunization. Further we found that storage of C57BL/6 RBCs makes them intrinsically immunogenic, rendering them capable of inducing DC maturation even in the absence of foreign epitopes. Therefore, RBCs become immunogenic under normal storage conditions, even in the absence of foreign epitopes, and activate immune responses through the adapter protein MyD88. These findings identify key immune signaling pathways contributing to RBC alloimmunization that could ultimately lead to identification of RBC-associated ligands necessary to initiate alloimmunization. Such an understanding is a critical step towards developing targeted therapies to mitigate RBC alloimmunization.

## Results

We utilized a transgenic mouse to model human blood transfusions in which RBCs express a triple fusion protein containing Hen egg lysozyme (HEL), Ovalbumin (OVA), and Duffyb (a human minor erythrocyte antigen), denoted the “HOD” mouse (Figure 1A) (14,32). This model allows us to dissect the specific receptors and pathways required for the generation of detrimental alloimmunity during transfusion. Storing HOD RBCs for transfusion allows for examination of the innate and adaptive immune response to a model human RBC antigen after transfusion of RBCs stored under conditions that are used in hospitals globally (13–25, 21). HOD RBCs processed and stored for 12 days at 4°C induced a robust anti-HOD alloantibody response post transfusion into wild type (WT) C57BL/6 mice. In contrast, the transfusion of fresh (non-stored) HOD blood induced minimal alloantibody production, at levels similar to un-transfused naïve mice (Figure 1B).

**Figure 1.**
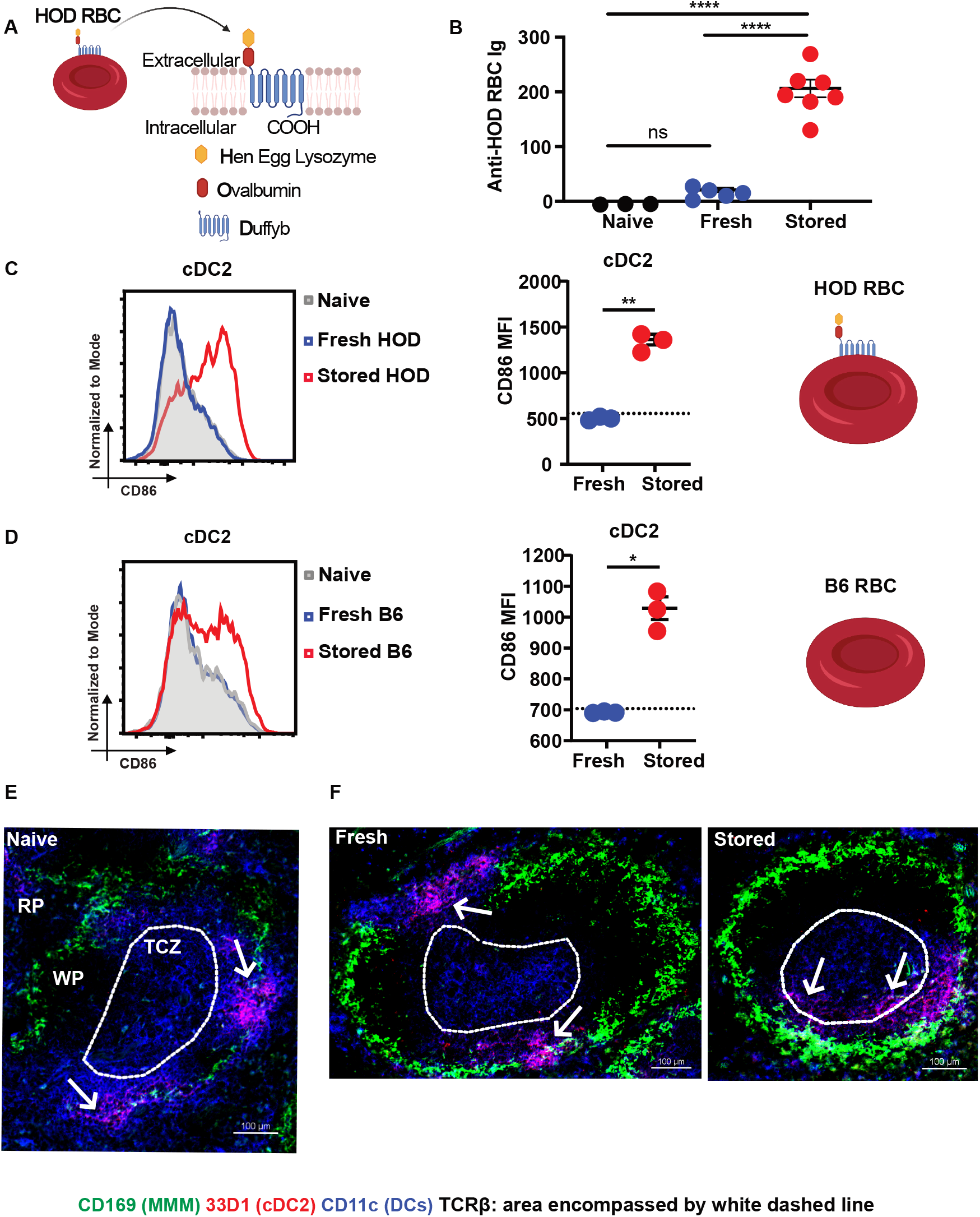
RBC storage but not alloantigen presence activates splenic cDC2s. (A) Model of HOD alloantigen containing Hen Egg Lysozyme (B cell epitope), Ovalbumin (T cell epitope), and Duffy_b_ (Human RBC antigen). Mice were transfused with freshly harvested RBCs or RBCs that had been stored for 12-14 days. (B) Anti-HOD RBC alloantibodies measured by flow cytometric crossmatch 21 days post fresh or stored RBC transfusion. (C) Flow cytometric analysis of splenic cDC2 CD86 expression 6h post fresh or stored HOD RBCs. Representative histogram of surface CD86 expression with accompanying quantification. Dashed line indicates value of naïve control. (D) Flow cytometric analysis of splenic cDC2 CD86 expression 6h post fresh or stored B6 RBCs without HOD alloantigen. Representative histogram of surface CD86 expression with accompanying quantification. Dashed line indicates value of naïve control. (E) Immunofluorescence (IF) microscopy of naïve spleen with arrows indicating localization of cDC2s in the bridging channel; (F) IF microscopy of spleen 6h post fresh or stored RBC transfusion with arrows indicating localization of cDC2s in the bridging channel after fresh transfusion and movement of cDC2s into and localization of cDC2s in the TCZ after stored RBC transfusion. TCZ = T-cell zone, RP = red pulp, WP = white pulp. Bar, 100 um. Error bars represent SEM for 3-6 mice per group. Groups were compared using Welch’s t-test. *P<0.05, **P<0.01, ****P<0.0001; ns, not significant. Representative data are shown for two to four independent experiments with n=3-7 mice per group.

Our prior work demonstrated that a subset of splenic conventional dendritic cells identified by staining with the antibody 33D1, known as type 2 conventional DCs (cDC2s), are required for T-dependent alloantibody production to stored RBCs (21). Once activated, these cDC2s upregulate the co-stimulatory molecule CD86 (33). We found that transfusion of stored, but not fresh HOD RBCs into C57BL/6 recipients induced CD86 expression on splenic cDC2s (Figure 1C, S1A). We next sought to determine whether DC activation by stored RBCs requires alloantigens on the RBCs. Similar to HOD RBCs, we found that syngeneic stored, but not fresh, C57BL/6 RBCs transfused into C57BL/6 recipients also induced CD86 expression on splenic cDC2s (Figure 1D), indicating that innate immune activation of cDC2s does not require alloantigen to be present on transfused RBCs.

In both mice and humans, the cellular architecture of the spleen is highly organized and dictates its function (34). cDC2s reside in specialized zones between the red and white pulp, called bridging channels, where they can readily sample the blood for antigens. Upon activation, cDC2s migrate into the T cell zone (TCZ) of the white pulp to provide stimulatory signals to CD4+ T cells (35). We found that transfusion of stored RBCs, regardless of alloantigen presence, resulted in cDC2 migration from bridging channels into the white pulp TCZ; this was not seen at steady state or in mice transfused with fresh RBCs (Figure 1E-F, S1B). Both DC migration into T cell zones and CD86 expression are necessary for subsequent T cell activation to occur, indicating that stored RBCs are capable of inducing requisite DC activation for adaptive immune responses, even in the absence of foreign antigen (33).

As our previous work had ruled out relevant NLRs as receptors for these mediators (13), we considered whether TLRs serve as receptors facilitating the immunogenicity of stored RBCs. All TLRs utilize TRIF and/or MyD88 as adapter proteins (31). We therefore used a *Trif^−/−^ Myd88^−/−^* mouse to determine whether TLRs play a role in the recognition of stored RBCs. In contrast to WT mice, *Trif^−/−^ Myd88^−/−^* mice had significantly decreased serum levels of pro-inflammatory cytokines and chemokines IL-6, KC (also known as CXCL1), and MCP-1 (also known as CCL2) two hours post transfusion (Figure 2A). Six hours post transfusion, *in vivo* splenic cDC2s from *Trif^−/−^ Myd88^−/−^* mice failed to upregulate CD86 in response to stored RBC transfusion (Figure 2B). Moreover, *Trif^−/−^ Myd88^−/−^* mice also demonstrated an impaired alloantibody response post transfusion, with anti-HOD antibody titers similar to those of naïve mice (Figure 2C).

**Figure 2.**
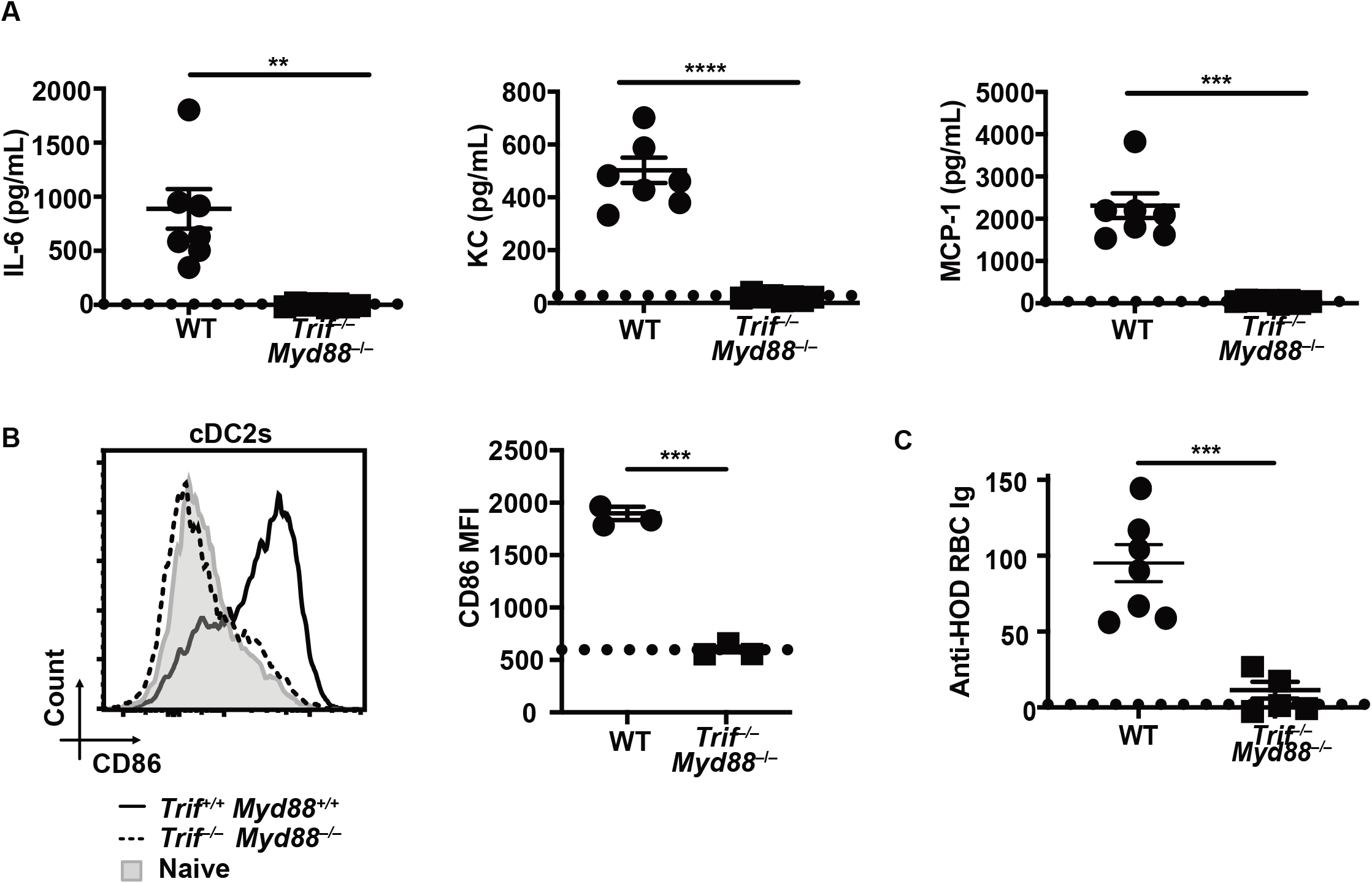
Loss of TLR signaling leads to impaired innate and adaptive immune responses to stored RBC transfusion. WT or *Trif^−/−^ Myd88^−/−^* mice were transfused with RBCs that had been stored for 12-14 days. (A) ELISA quantification of IL-6 (left), KC (middle), or MCP-1 (right) in recipient serum 2h post transfusion. Dashed line indicates value of naïve control. (B) Flow cytometric analysis of splenic cDC2 CD86 expression 6h post stored RBC transfusion into WT or *Trif^−/−^ Myd88^−/−^* mice. Representative histogram of surface CD86 expression with accompanying quantification. (C) Anti-HOD RBC alloantibodies measured by flow cytometric crossmatch 21 days post stored RBC transfusion. Dashed line indicates value of naïve control. Error bars represent SEM for n=3-7 mice per group. Groups were compared using Welch’s t-test. **P<0.01, ***P<0.001, ****P<0.0001.

We next sought to differentiate the effects of MyD88 and TRIF deficiency independe *Myd88^−/−^* or *Trif^−/−^* mice. Different PRRs signal through unique downstream pathways utilizing distinct adaptor proteins (27, 36). Through the use of *Trif^−/−^* and *Myd88^−/−^* mice, we aimed to narrow down the number of candidate receptors that stored RBC products can stimulate. MyD88 is a pleiotropic adaptor molecule and plays an important role in both innate and adaptive immune signaling pathways (37–42). Unlike WT mice, *Myd88^−/−^* mice failed to produce the pro-inflammatory chemokine MCP-1 two hours following stored RBC transfusion (Figure 3A). However, *Trif^−/−^* mice secreted MCP-1 at levels similar to WT mice (Figure 3B). This suggests that MyD88, but not TRIF, plays an important role in the early innate immune response to stored RBC products. Notably, *Myd88^−/−^* mice were also unable to mount an alloantibody response to HOD RBCs post RBC transfusion (Figure 3C). In contrast, *Trif^−/−^* mice demonstrated anti-HOD alloantibody levels comparable to WT mice (Figure 3D). Therefore, receptors that require signaling through the adaptor protein TRIF to function, such as TLR3, are not required for the alloimmune response to stored donor HOD RBCs.

**Figure 3.**
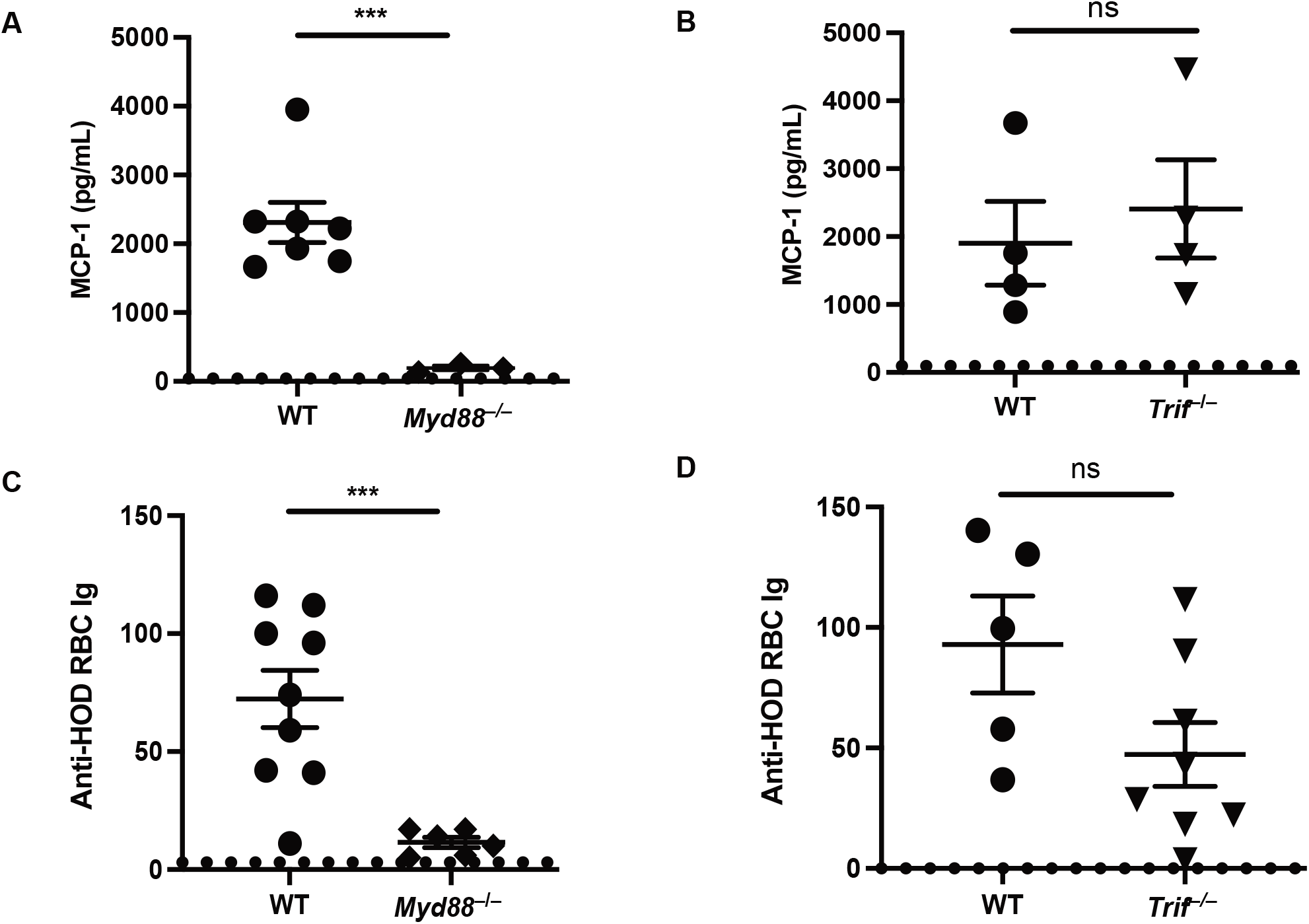
MyD88 but not TRIF is required for the innate and adaptive response to stored RBC transfusion. *Myd88^−/−^*,*Trif^−/−^*, or WT mice were transfused with HOD RBCs that had been stored for 12-14 days. (A) ELISA quantification of MCP-1 in *Myd88^−/−^* recipient serum 2h post transfusion. Dashed line indicates value of naïve control. (B) ELISA quantification of MCP-1 in *Trif^−/−^* recipient serum 2h post transfusion. Dashed line indicates value of naïve control. (C) Anti-HOD RBC alloantibodies in *Myd88^−/−^* recipients measured by flow cytometric crossmatch 21 days post stored RBC transfusion. Dashed line indicates value of naïve control. (D) Anti-HOD RBC alloantibodies in *Trif^−/−^* recipients measured by flow cytometric crossmatch 21 days post stored RBC transfusion. Dashed line indicates value of naïve control. Error bars represent SEM for n=3-9 mice per group. Groups were compared using Welch’s t-test. ***P<0.001; ns, not significant.

We next addressed which cell type requires MyD88 signaling for RBC-induced activation. The spleen is responsible for filtration of the blood and monitoring of the circulation for antigens (34). To determine responsiveness to stored RBCs, we examined upregulation of the activation marker CD86 on different splenic immune cell populations *in vivo* six hours following transfusion. Using this readout, stored RBCs did not directly activate follicular B cells or marginal zone B cells (Figure S2A-B). We also examined splenic resident innate immune cells including cDC subsets, macrophages, monocytes, and plasmacytoid DCs. Consistent with our previous findings both subsets of cDCs were strongly activated in response to stored transfusion (Figure 4A). Upregulation of CD86 by cDCs was completely abrogated in *Myd88^−/−^* mice, indicating that functional MyD88 signaling is required for cDC activation to stored RBCs (Figure 4A). In contrast, monocytes did not become activated in response to stored RBCs (Figure 4A). Although pDC and macrophage CD86 upregulation in response to RBC transfusion was statistically significant between control and knockout mice, it might not be biologically significant given the low level of activation relative to cDCs (Figure 4A, S2A). Our previous work demonstrated that cDC2s are the primary DC subset responsible for alloantibody production in response to RBC transfusion in part by selectively migrating to the CD4+ T cell region of the white pulp (21, 35). We found that cDC2 migration into the white pulp of the spleen is impaired in *Myd88^−/−^* mice six hours after stored RBC transfusion relative to spleens from control mice (Figure 4B). Taken together, these data show that stored RBCs activate cDCs through a MyD88 dependent pathway.

**Figure 4.**
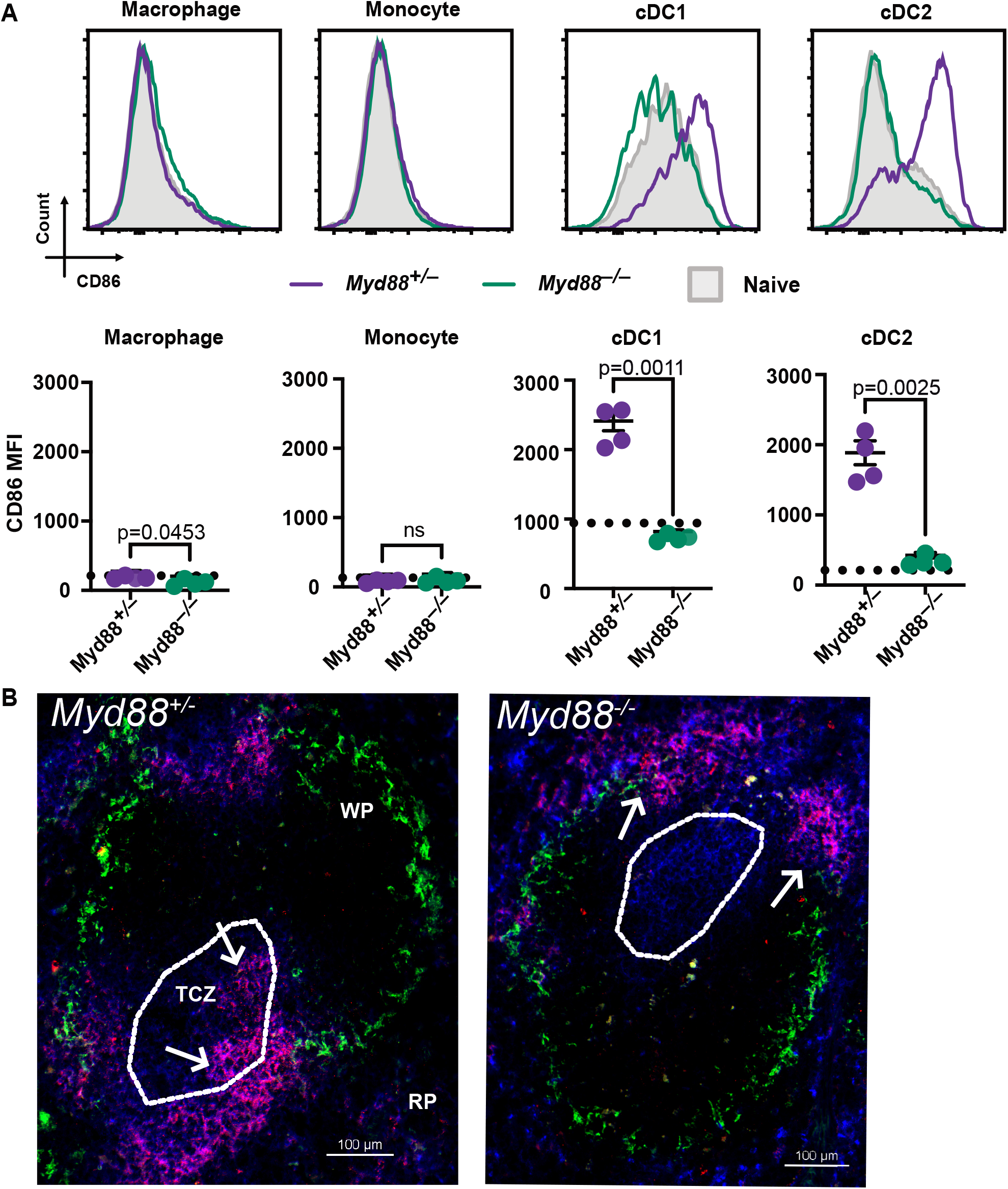
Stored RBC transfusion activates conventional DCs through a MyD88 dependent signaling pathway. *Myd88^+/–^* or *Myd88^−/−^* mice were transfused with C57BL/6 RBCs that had been stored for 12-14 days. (A) Flow cytometric analysis of CD86 on splenic macrophage, monocyte, cDC1, and cDC2 (gating shown in Supplement 1A) 6h post stored RBC transfusion into *Myd88^+/–^* (purple) or *Myd88^−/−^* (green) mice. Representative histogram of surface CD86 expression (top) with accompanying quantification (bottom). Dashed line indicates value of naïve control. Error bars represent SEM for n=4 mice per group. *Myd88^+/–^* or *Myd88^−/−^* groups were compared using Welch’s t-test. ns, not significant. (B) Immunofluorescent microscopy of *Myd88^+/–^* or *Myd88^−/−^* spleen 6h post stored RBC transfusion with arrows indicating localization of cDC2s in the bridging channel in *Myd88^−/−^* and movement of cDC2s into and localization of cDC2s in the TCZ in *Myd88^+/–^*. Bar, 100 um.

## Discussion

In this study, we have demonstrated that MyD88 is required for the innate (DC activation) and adaptive (alloantibody production) immune response to transfused stored RBCs, suggesting that one or more MyD88-dependent TLRs are responsible for sensing transfused RBCs after storage. In contrast, TRIF was not required for alloantibody production, indicating TLR3 is unlikely to be required in recognizing transfused RBCs. Further we found that storage of syngeneic (i.e., non-allogeneic) C57BL/6 RBCs also renders the RBCs able to induce DC maturation in the absence of foreign antigens. This demonstrates that alloantigen is not required for innate immune stimulation by RBCs. Future work aiming to identify whether such an effect also occurs during storage of human RBCs will be important. Interestingly, some blood donors demonstrate better RBC storage characteristics than others; in parallel, some transfusion recipients are considered “responders” whereas others are “non-responders”, in terms of alloantibody production. More work is needed to identify the immunologic underpinnings of these donor and recipient effects on alloimmunization. We posit that defining the innate immune pathways that regulate the alloimmune response is the first step towards understanding and ultimately preventing RBC alloimmunization.

Future work is needed to identify whether MyD88 is acting in a DC-intrinsic versus -extrinsic manner; this will provide mechanistic insights into the process of RBC alloimmunization and might help narrow down candidate receptors (e.g., particular TLRs) through which RBC alloimmunization is initiated. Although we have demonstrated that cDCs are activated during the course of transfusion, it is possible that additional accessory cells in the spleen, such as the multiple macrophage populations in and around the bridging channel, are activated through MyD88 to provide DC activating signals. We have previously observed that macrophage-DC crosstalk in the spleen impacts adaptive immunity to bacteria (30); the contribution of such crosstalk in the adaptive immune response to stored RBCs requires further investigation. Other cells in the spleen might also influence DC-dependent T cell priming to allogenic RBCS. We observed limited pDC activation in response to stored RBC transfusion and the role of such cells in priming naïve T cells is controversial (43); however, pDCs have been shown to increase transfused RBC consumption during states of inflammation (44). While no difference in marginal zone B cell activation was observed following RBC transfusion, recent studies demonstrate that RBC alloimmunization likewise requires marginal zone B cells, suggesting that cDCs may work in concert with marginal zone B cells to facilitate alloimmunization (45–47).

Understanding which innate immune receptors are responsible for the initiation of the RBC alloimmune response will help identify possible immunogenic compounds released by or bound to RBCs that enhance immunogenicity and therefore risk for alloantibody production. Ultimately, this information could be useful for developing prophylactic strategies to either block receptor signaling or alter RBC units to prevent RBCs alloimmunization.

## Materials and Methods

### Mice

WT C57BL/6 mice were purchased from Charles River. HOD mice were generated as previously described ^32^. *Trif*^−/−^ mice were purchased from The Jackson Laboratory. *Myd88*^−/−^ mice were purchased from The Jackson Laboratory and bred in our facility to generate *Myd88*^−/−^ and *Myd88*^+/–^ littermate controls to facilitate breeding, to create MyD88 homozygous and heterozygous littermate controls, and to control for microbiota driven effects. *Trif* ^−/−^ *Myd88*^−/−^ mice were generated by breeding mice purchased from The Jackson Laboratory in our facility.

### RBC Transfusion Model

Red blood cells (RBCs) were collected from HOD or C57BL/6 mice into a 13% Citrate Phosphate Dextrose Adenine anti-coagulant (final volume) and leukoreduced using a murine adapted Pall Acrodisc PSF 25mm WBC filter with Leukosorb Media. Leukoreduced blood was adjusted to a hematocrit of approximately 75% as previously described ^14,25^. RBCs were then transfused immediately or stored for 12-14 days at 4°C ^21^. 100uL of RBCs was transfused retro-orbitally into recipient mice.

### Enzyme-linked Immunosorbent Assay (ELISA)

Serum samples were analyzed by ELISA for MCP-1, KC, or IL-6. MCP-1, KC, and IL-6 were detected with respective mouse cytokine ELISA kits (R&D) according to enclosed protocol.

### Flow cytometry analyses

Spleens were manually digested, washed in 2% FBS in PBS, and incubated with fluorescent antibodies for 30 minutes at 4°C. Cells were analyzed on either a Cytoflex (Beckman Coulter) or MACSQuant (Miltenyi) flow cytometer and analyzed using FlowJo software (Tree Star). The following antibodies were used for staining different cell subsets (Biolegend unless otherwise noted): F4/80 (BM8), CD11c (N418), CD86 (GL 1), B220 (RA3-6B2), Ly6C (HK1.4), CD8a (53-6.7), TCRb (H57-579), CD11b (M1/70), MHC II (M5/114.15.2), CD23 (B3B4), CD19 (6D5), CD21/35 (7E9), NK1.1 (PK136), 33D1 (33D1), CD169 (3D6.112), and goat anti-mouse Ig (BD Biosciences Cat #550826).

### Immunofluorescence microscopy

Spleens were harvested and then dehydrated for an hour in successive solutions of 10, 20, and 30 percent sucrose. Spleens were mounted in a cryomold with O.C.T. Compound (Tissue-Tek, Sakura) and frozen at −80°C before sectioning into 7 micron slices. All images were acquired with the Nikon eclipse Ti scope using 10x objectives.

### Statistics

All statistical analyses were performed using GraphPad Prism software. Data were analyzed with Welch’s unpaired t-test or one-way ANOVA with Tukey test.

### Serum analysis

Transfused mice were terminally bled via cardiac puncture or by retro-orbital bleed 21 days post RBC transfusion for anti-HOD alloantibody measurements, or 6 hours post transfusion for acute serum cytokine measurements. 21 days is the peak of the alloantibody response based on our previous work ^21^. Blood was left to clot, the clot was removed, and then the remaining blood was centrifuged at 1,500 x g for 10 minutes. The liquid serum was pipetted off and frozen at −20°C. To identify anti-HOD alloantibodies, sera were plated in a 96-well U bottom plate, incubated with 3uL of HOD RBCs for 30 minutes, washed, and incubated with APC anti-Ig antibodies for 30 more minutes. Sera were washed again and then run on the cytometer to detect total anti-HOD Igs. Anti-RBC antibodies in figures are all anti-HOD total Igs and were quantified by mean fluorescence intensity (MFI) in arbitrary units.

### Study approval

All protocols were approved by the Institutional Animal Care and Use Committee of the Yale Animal Resource Center.

## Author Contributions

AS and LRH wrote the manuscript. AS, LRH, LX, SC, SML, AEG, and DL conducted experiments, acquired data, and analyzed data. MS and JEH provided reagents and mice. KEH, SRS, CJL, JCZ, JEH, and SCE designed research studies. All authors reviewed the manuscript. AS is listed first with the understanding that LRH will be listed first on the subsequent paper.

## Acknowledgements

We would like to thank Jelena Medved, Mark Firla and Joan Goldstein for technical assistance. We would also like to thank Jennifer S. Chen, Sam Olyha, Kenneth Zhou, and Emily Siniscalco for their discussion and review of the manuscript.

---

**Supplement 1 (for Figure 1).**
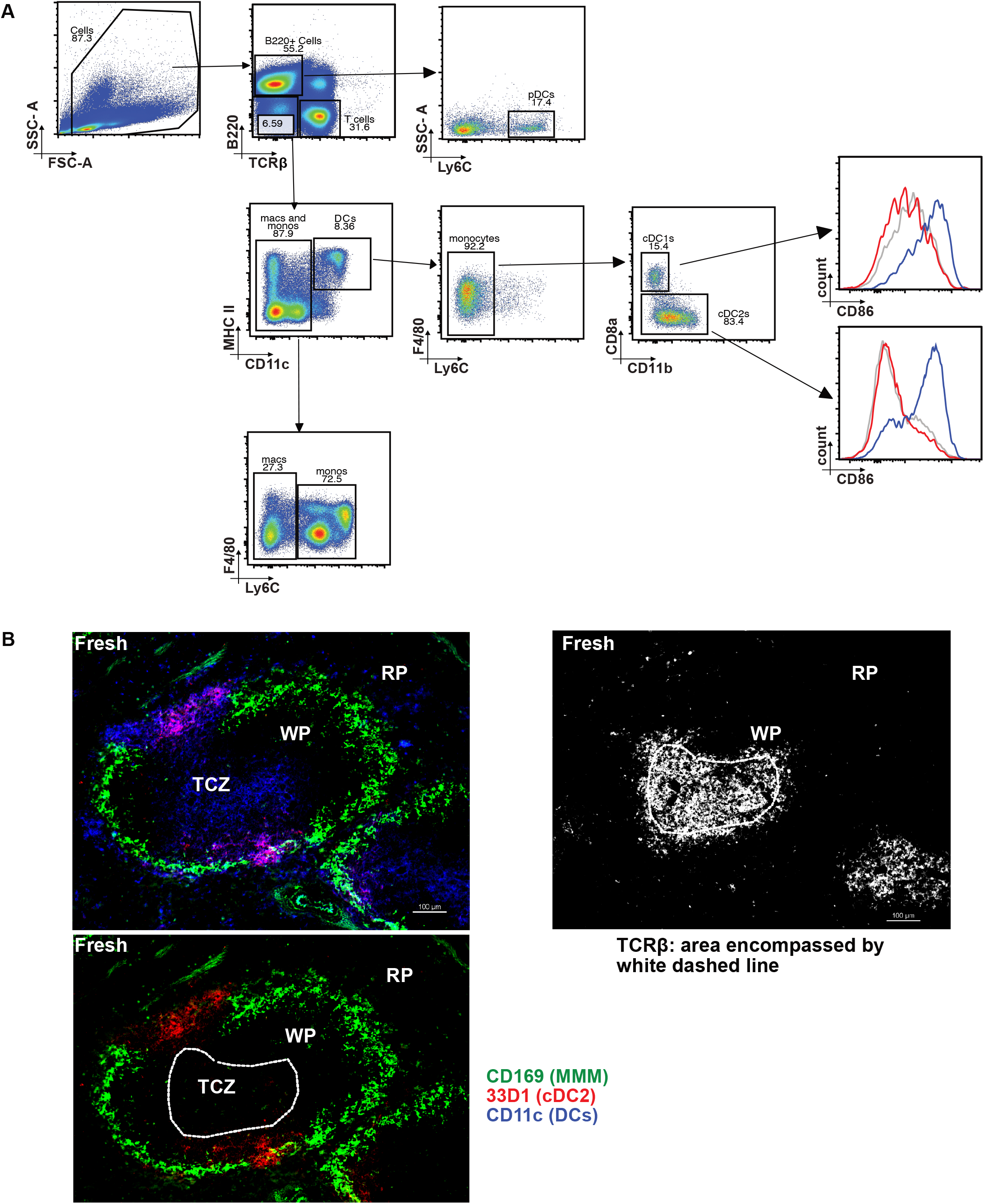
(A) Flow gating strategy for pDC, macrophage, monocyte, cDC1, and cDC2 cells. (B) Image processing strategy for IF in 1E and F; top left: 3 color composite image without T-cell zone area circled, top right: strategy for defining T-cell zone, bottom left: 33D1 (cDC2) and CD169 (marginal zone metallophilic macrophage) staining only. Bar, 100 um.

**Supplement 2 (for Figure 4).**
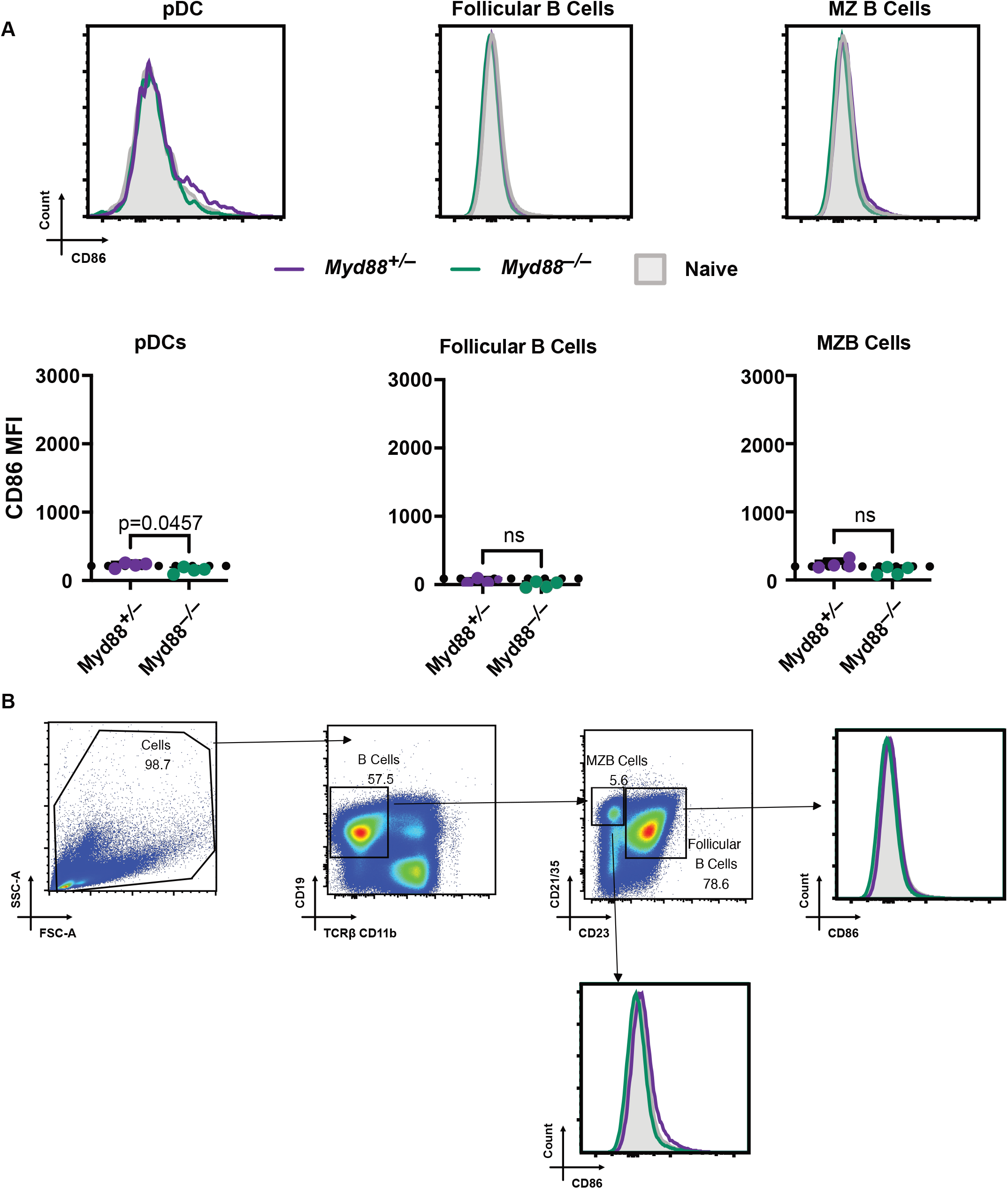
*Myd88^+/–^* or *Myd88^−/−^* mice were transfused with RBCs that had been stored for 12-14 days. (A) Flow cytometric analysis of splenic pDC, Follicular B cell, and MZB cell CD86 expression 6h post stored RBC transfusion into *Myd88^+/–^* or *Myd88^−/−^* mice. Representative histogram of surface CD86 expression with accompanying quantification. Dashed line indicates value of naïve control. (B) Flow gating strategy for Follicular B cells and MZB cells. Error bars represent SEM for n=4 mice per group. Groups were compared using Welch’s t-test. ns, not significant.

